# Fluorescence cross-correlation spectroscopy quantifies affinity, cooperativity, and kinetic stability in ternary protein complexes

**DOI:** 10.64898/2026.07.16.738902

**Authors:** Stefan H. Mueller, Gayathri Mohanan, Mahmoud A. S. Abdelhamid, Christopher P. Toseland, Timothy D. Craggs

**Author notes:** Corresponding authors: Stefan H. Mueller, Timothy D. Craggs. **Competing Interest Statement:** S.H.M., G.M., M.A.S.A., C.P.T. and T.D.C. are employees and/or shareholders in Exciting Instruments Ltd.

## Abstract

Many biological processes and emerging therapeutic modalities rely on higher-order protein complexes whose properties cannot be predicted from their constituent binary interactions. However, methods for directly quantifying affinity, cooperativity, and kinetic stability within such assemblies remain limited. Here, we establish fluorescence cross-correlation spectroscopy (FCCS) as a solution-phase approach for characterizing multicomponent protein interactions and apply it to the clinically important HER2-targeting antibodies trastuzumab and pertuzumab.

Using fluorescently labelled HER2, trastuzumab, and pertuzumab, we quantified binary binding affinities, directly measured ternary complex formation, and characterized the dissociation kinetics of binary and ternary complexes. FCCS measurements revealed positive cooperativity in the formation of the HER2–trastuzumab–pertuzumab ternary complex, while dissociation experiments demonstrated that the ternary complex is kinetically more stable than the corresponding binary interactions. Together, these findings provide direct solution-phase evidence that cooperative interactions stabilize the HER2–trastuzumab–pertuzumab complex, offering a molecular explanation for the enhanced efficacy of dual HER2 targeting in cancer therapy. More broadly, this work demonstrates that FCCS can robustly quantify affinity, cooperativity, and kinetic stability of multicomponent protein complexes using a commercially available platform. We provide a broadly accessible framework for studying higher-order protein interactions and supporting the development of next-generation multispecific and combination therapeutics.

**Significance Statement:** Many proteins function as part of multicomponent complexes, yet most experimental methods characterize interactions only one pair at a time, or indirectly through secondary reporters, or necessitating saturation of binary interactions first. We show that fluorescence cross-correlation spectroscopy can directly quantify how multiple binding partners interact simultaneously by measuring affinity, cooperativity, and kinetic stability in solution. Applying this approach to the clinically important HER2-targeting antibodies trastuzumab and pertuzumab reveals cooperative stabilization of their ternary complex. Because the measurements are performed on a commercially available instrument and require no surface immobilization, this broadly accessible method should facilitate mechanistic studies of complex biomolecular interactions and aid the development of multispecific and combination therapeutics.

## Introduction

Biological function emerges from networks of interacting proteins that assemble into dynamic molecular complexes (1, 2). Processes as diverse as signal transduction, immune recognition, transcriptional regulation, and intracellular trafficking depend on the formation and dissociation of higher-order protein assemblies(3, 4). The composition, stability, and dynamics of these complexes determine how biological information is transmitted and regulated, making quantitative characterization of multi-component protein interactions a central goal of modern molecular biology.

The ability to manipulate protein complexes has likewise become a major objective in therapeutic development. Examples include molecular glues and PROTACs that induce neomorphic interactions between E3-ligases and disease-related proteins (5, 6), bispecific antibodies that physically bridge distinct cellular targets (7), and multispecific biologics that exploit simultaneous engagement of multiple epitopes to enhance functional activity (8, 9). In such systems, drug efficacy often depends not only on binding affinity but also on cooperative interactions that influence the formation and stability of higher-order assemblies (10–13).

Despite their importance, quantitative characterization of higher-order protein assemblies remains challenging. Structural methods provide detailed molecular architecture but offer limited insight into equilibrium populations and dissociation dynamics in solution, as well as diverse conformational dynamics (14, 15). Surface-based biophysical techniques such as surface plasmon resonance and biolayer interferometry have transformed the study of binary interactions yet become increasingly difficult to execute and interpret for multi-component systems where several molecular species coexist simultaneously (11, 16, 17). In particular, directly resolving specific ternary and higher-order complexes and determining how binding of one component influences another remains a major experimental challenge (18, 19).

The human epidermal growth factor receptor 2 (HER2) is involved in a variety of signaling pathways leading to cell proliferation. It plays a key role in diagnostics and therapy for up to 30% of breast cancers and gastric/gastroesophageal cancers(20).The HER2-targeting antibodies trastuzumab and pertuzumab represent one of the most successful examples of combination antibody therapy. Trastuzumab, first approved in 1998 for HER2-positive breast cancer, transformed outcomes for patients with HER2-driven disease (21). The subsequent approval of pertuzumab in combination with trastuzumab in 2012 further improved clinical efficacy and established dual HER2 blockade as a standard therapeutic strategy (22, 23). Because the two antibodies bind distinct epitopes and were originally proposed to act through complementary mechanisms — including inhibition of HER2 signaling by trastuzumab and interference with HER2 dimerization by pertuzumab — the magnitude of the clinical benefit stimulated considerable interest in whether direct molecular synergy contributes to their combined activity (24–27). This question has remained unresolved. Structural studies have demonstrated simultaneous binding of both antibodies to HER2 and generally support independent occupancy of their respective epitopes (18). Likewise, several biophysical studies have reported little evidence for cooperative binding (28, 29). In contrast, computational analyses and studies probing HER2 conformational dynamics have suggested that binding of one antibody may influence the interaction of the other through allosteric effects (12, 30–32). Together, these observations have produced a conflicting picture of how the trastuzumab–pertuzumab–HER2 ternary complex is assembled and stabilized.

Here we establish fluorescence cross-correlation spectroscopy (FCCS) as a quantitative platform for characterizing higher-order protein complexes directly in solution. FCCS enables direct observation of specific molecular assemblies through co-diffusion of spectrally distinct binding partners, allowing equilibrium complex formation and dissociation kinetics to be measured within a unified microplate-compatible workflow. Applying this approach to the trastuzumab–pertuzumab– HER2 system, we directly quantify formation of the ternary complex and measure its equilibrium and kinetic properties in solution. While ternary complex assembly is broadly consistent with simultaneous binding to distinct HER2 epitopes, quantitative analysis reveals positive cooperativity in both equilibrium complex formation and dissociation kinetics. These findings help reconcile conflicting observations in the literature by suggesting that cooperative stabilization of the trastuzumab–HER2–pertuzumab complex is present but may be difficult to resolve in methods that do not selectively distinguish ternary complexes from the underlying heterogenous mixture of binary and higher-order states. More broadly, our results demonstrate how direct observation of higher-order protein assemblies can uncover mechanistic features of biomolecular interactions that remain obscured in conventional structural and ensemble-averaged biophysical measurements.

## Results

### Characterization of Trastuzumab- and Pertuzumab-HER2 Affinities in Solution

To establish a solution-phase assay for quantifying antibody-antigen interactions or protein-protein interactions more generally by FCCS, we first characterized the interactions of trastuzumab and pertuzumab with the extracellular domain of HER2.

We labelled full-length antibodies and the extracellular domain of HER with spectrally separated fluorescent dyes using NHS-chemistry, achieving degrees of labeling between 0.8-1.8 (HER2) and 2.8-5.2 (antibodies, see supplementary figure and table S1). As molecules enter and leave the observation volume, fluorescence fluctuations are recorded independently in the HER2 and antibody detection channels. Unbound molecules contribute only to their respective channels and auto-correlation functions. Binding between labelled HER2 and labelled antibodies results in co-diffusion of the two fluorescent species through the confocal volume and this produces fluctuations detectable simultaneously in both channels, giving rise to a cross-correlation signal (Figure 1A). The cross-correlation does not rely on changes in diffusion or molecular mass of the complex, and, unlike fluorescence resonance energy transfer (FRET) measurements, is independent of the molecular geometry of the formed complex. Because autocorrelation amplitudes are inversely proportional to the number of fluorescent molecules in the observation volume, increasing HER2 concentration leads to a progressive decrease in the HER2 autocorrelation amplitude. The cross-correlation amplitude, however, is proportional to the number of co-diffusing complexes divided by the product of the total particle numbers detected in each of the two detection channels. Furthermore, optical artifacts, such as imperfect overlap of the confocal volumes in two colors as well as potential FRET can make it difficult to extract absolute concentrations of complexes. To account for these effects, the cross-correlation amplitude was normalized to the HER2 auto-correlation amplitude. Since the antibody concentration was held constant throughout the titration, the resulting normalized cross-correlation signal is proportional to the fraction of HER2 molecules participating in antibody–HER2 complexes. As HER2 concentration increases, the normalized cross-correlation signal therefore reports the relative increase in complex formation. The normalized cross-correlation ratio serves as a robust ratiometric measure of binding, independent of signal fluctuations caused by other factors.

**Figure 1.**
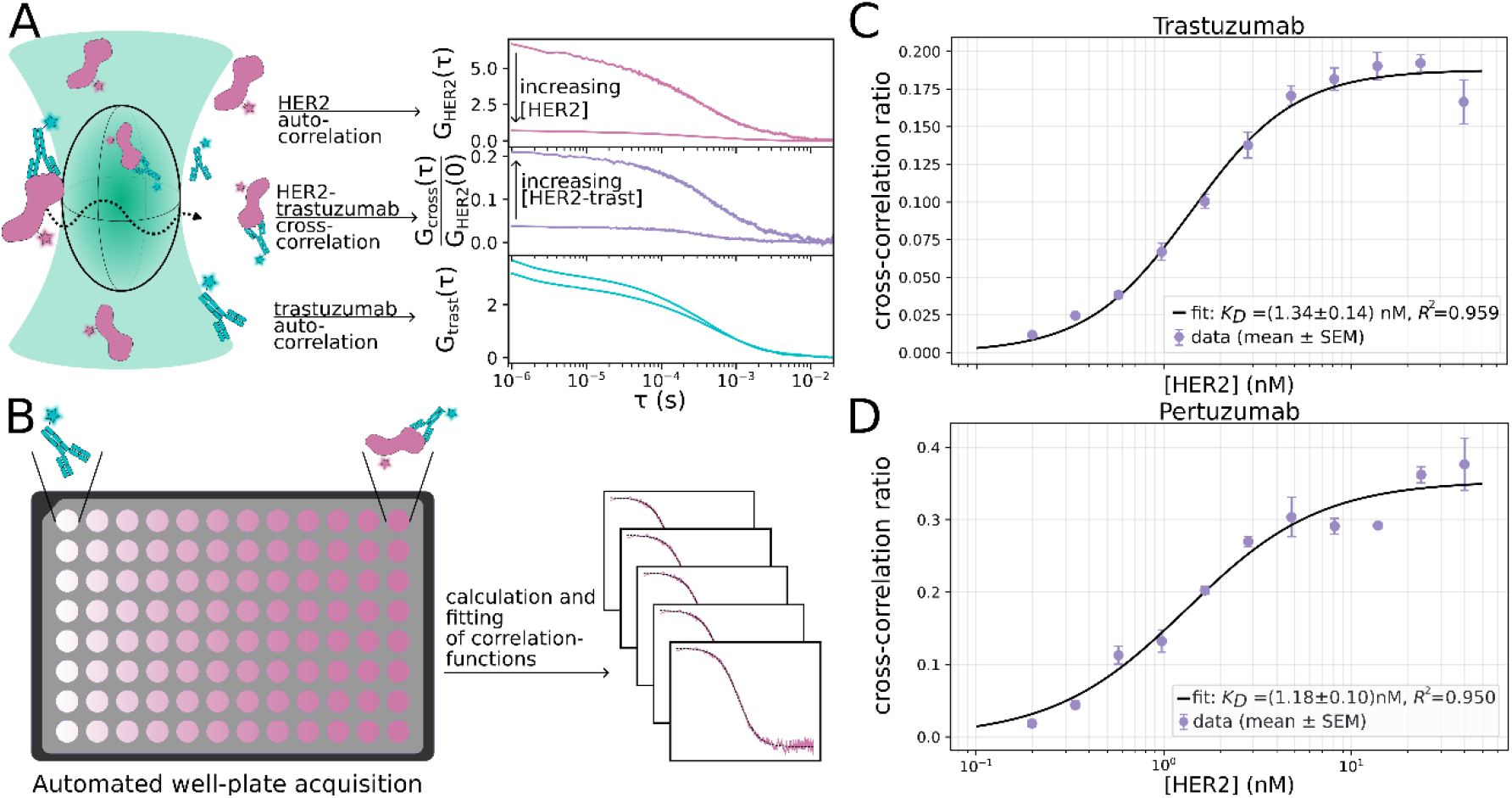
(**A**) left: schematic of HER2 (magenta)/trastuzumab (cyan) (co-)diffusing through the confocal volume and the impact of binding on auto-correlation functions (magenta/cyan) as well as cross-correlation function (purple). **(B)** Schematic of multiplexed FCCS-measurements in 96-well plate format. (**C)** Titration of HER2 to trastuzumab. Error bars depict the standard error of the mean. The data is fit with a quadratic binding model (see methods). (**D**) Titration of HER2 to pertuzumab. Error bars depict the standard error of the mean. The data is fit with a Hill equation (see methods)

Equilibrium titrations were performed in a multiplexed 96-well format confocal fluorescence spectrometer (Figure 1B). The measured and normalized cross-correlation amplitudes were corrected for spectral bleed-through and background (see methods). The resulting data were fit with quadratic ligand-binding equations. Dissociation constants for trastuzumab and pertuzumab to HER2 were found be *K*_d,trast_ = 1.34 ± 0.14 nM and *K*_d,pert_ = 1.18 ± 0.10 nM (mean ± SEM, n=3, Figure 1C/D). Within experimental uncertainty, both antibodies exhibited comparable affinities for HER2 under the conditions used here.

### Competitive-chase FCCS Confirms Independent Binding of Trastuzumab and Pertuzumab to Distinct Epitopes on HER2

We next investigated whether FCCS could resolve dissociation kinetics while simultaneously probing the effects of binding a second antibody to preformed antibody–HER2 complexes. Binary trastuzumab–HER2 and pertuzumab–HER2 complexes were preformed at 10 nM antibody and HER2 (approximately tenfold above the measured equilibrium dissociation constants) and subsequently challenged with a 30-fold excess of the corresponding unlabeled antibody. Dissociation was monitored through the loss of normalized cross-correlation over time (Figure 2A).

**Figure 2.**
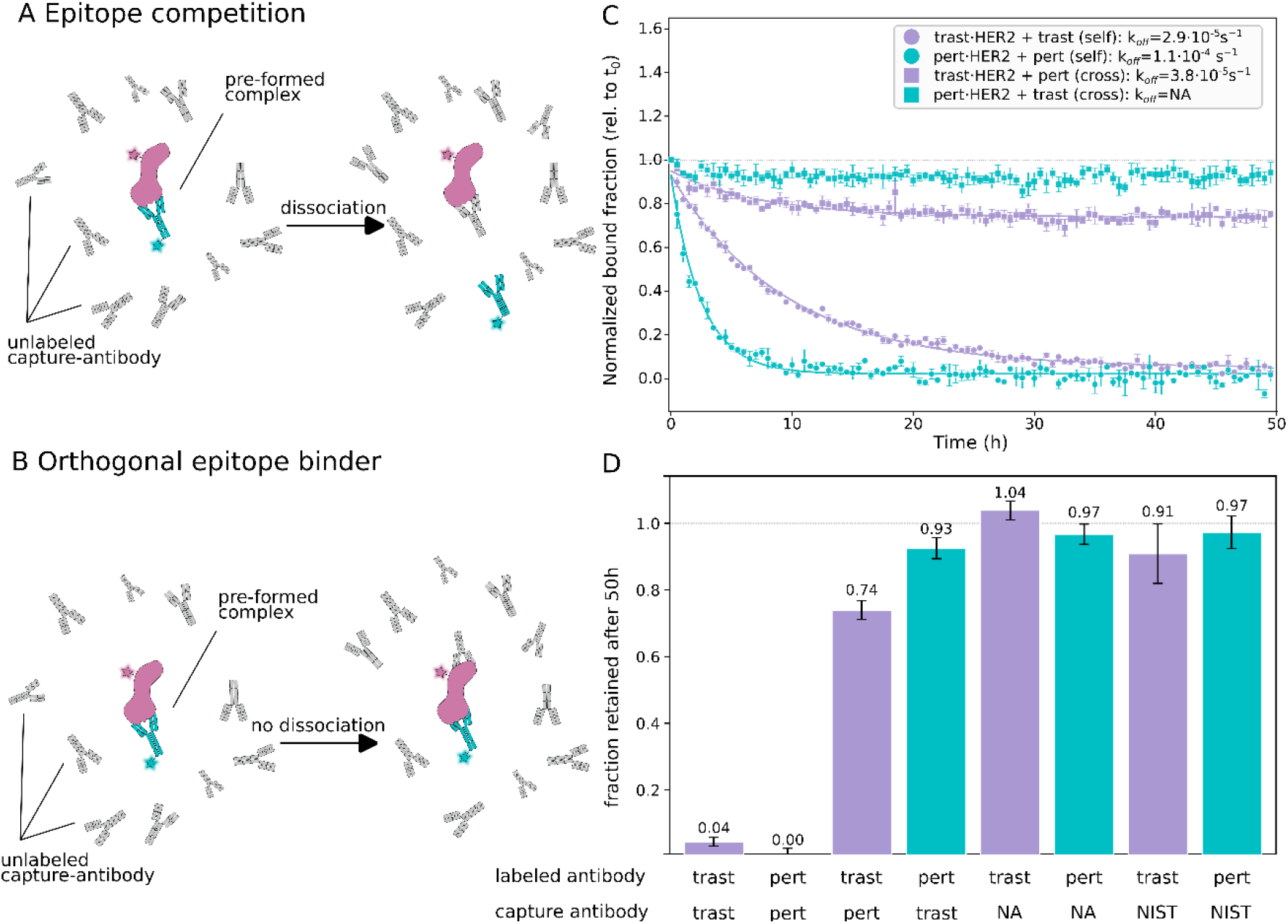
(**A, B**) Schematic of epitope competition (top)/orthogonal binder (bottom) experiment: Preformed complexes of labelled antibodies and targets are subjected to large excesses of unlabeled competitors. Replacement of the labeled antibodies by unlabeled competitors results in loss of signal, orthogonal binders leave the signal unchanged. (**C)** Dissociation of trastuzumab (purple) and pertuzumab (cyan). The controls show trastuzumab competing with unlabeled pertuzumab and vice versa. (**D**) Normalized cross-correlation fraction retained after 48 h in presence of capture antibodies trastuzumab (trast), pertuzumab (pert), None (NA) or non-specific control IgG (NIST).

Addition of the corresponding unlabeled antibody resulted in a gradual loss of cross-correlation for both trastuzumab–HER2 and pertuzumab–HER2 complexes, reflecting replacement of the labelled antibody following spontaneous dissociation. Trastuzumab dissociated substantially more slowly than pertuzumab under identical conditions (Figure 2B). In contrast, challenging HER2-bound trastuzumab with unlabeled pertuzumab and vice versa revealed an asymmetric response (Figure 2C,D). Addition of excess unlabeled trastuzumab to preformed pertuzumab–HER2 complexes produced no measurable reduction in the cross-correlation signal, consistent with independent binding of separate epitopes. Surprisingly, addition of excess unlabeled pertuzumab to preformed trastuzumab–HER2 complexes resulted in a small but reproducible decrease in the cross-correlation signal, corresponding to an approximately 25% reduction in labelled complexes. This suggests that pertuzumab binding perturbs a subset of trastuzumab–HER2 complexes. Challenge with an unrelated control antibody (NIST IgG1) as well as the absence of any unlabeled antibody produced no measurable reduction in cross-correlation for either complex, excluding non-specific IgG-mediated effects (Figure 2D and supplementary Figure S6).

Single-exponential fitting of the dissociation trajectories yielded molecular dissociation rates of (2.9 ± 0.2) × 10^-5^s^-1^ for trastuzumab and (1.1 ± 0.5) × 10^-4^s^-1^ for pertuzumab, confirming the slower dissociation of trastuzumab. Control reactions in the absence of challenge remained stable over the full 48 h observation period, demonstrating that the observed signal losses arose from antibody exchange rather than spontaneous decay or instrumental drift. Together, these results demonstrate that FCCS enables direct measurement of high-affinity antibody dissociation kinetics while simultaneously revealing subtle, epitope-specific interactions between antibodies binding distinct sites on the same target protein.

### Direct Interrogation of the Trastuzumab-Her2-Pertuzumab Ternary Complex

Finally, we aimed to characterize the formation of the ternary complex of trastuzumab-HER2-pertuzumab, previously hypothesized to be critical for the synergy of treatment with both antibodies. Previous studies have shown that trastuzumab and pertuzumab bind distinct epitopes on HER2 (Figure 3C). Here, using fluorescently labeled trastuzumab and pertuzumab in combination with unlabeled HER2, the cross-correlation signal is proportional to the interactions of both trastuzumab and pertuzumab. We find this interaction to be negligible in the absence of HER2 (supplementary Figure S4,S5). In the presence of HER2 the cross-correlation, therefore, provides a direct measure for the formation of the ternary complex of trastuzumab-HER2-pertuzumab.

**Figure 3.**
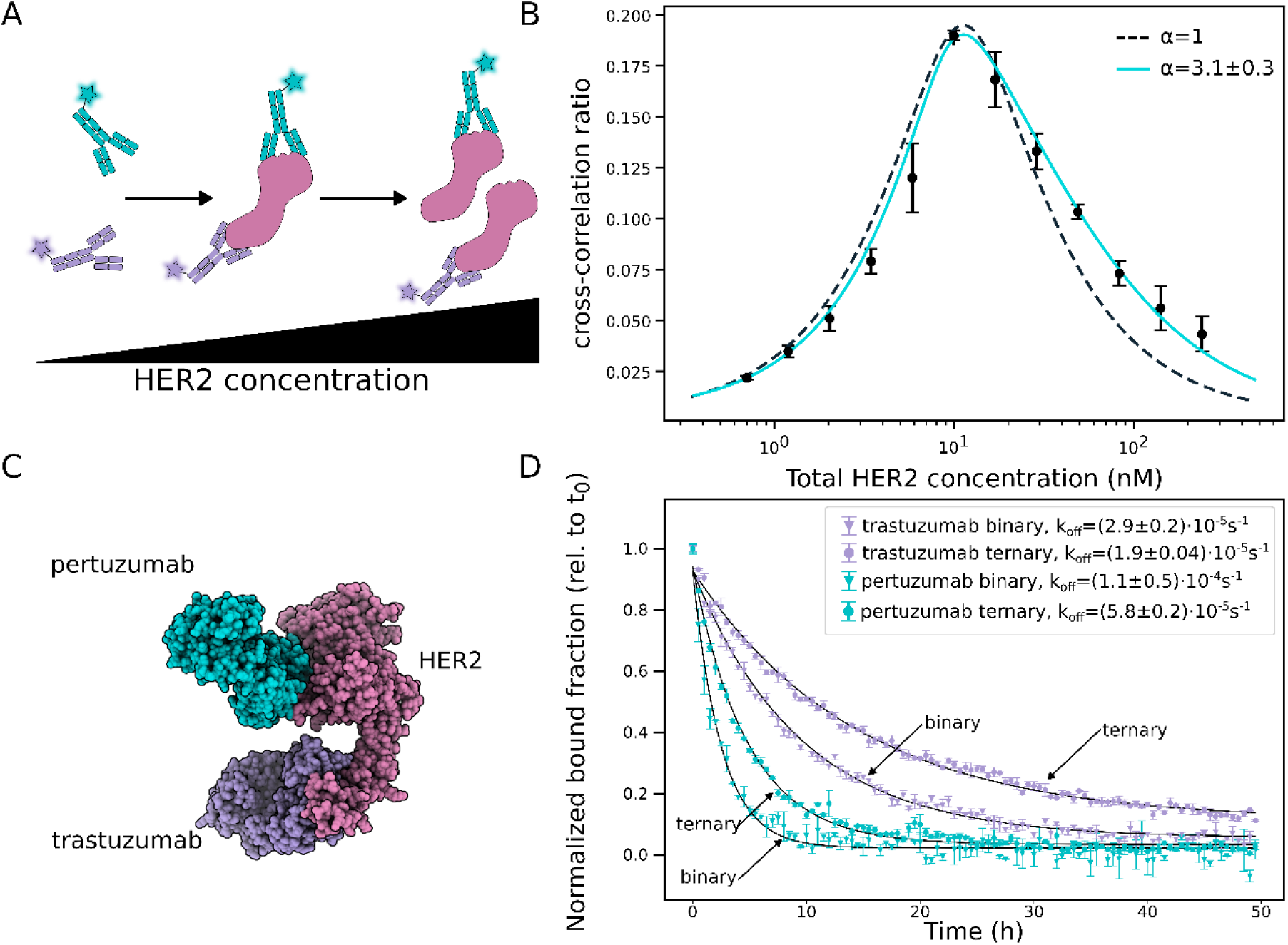
**(A)** Schematic of ternary complex formation: As HER2 concentration increases more ternary complex is formed. In large excess of HER2 the antibodies are sequestered and rarely bound to the same molecule of HER2, leading to the hook-effect. **(B)** Ternary complex formation of Trastuzumab-HER2-Pertuzumab as a function of unlabeled Her2 concentration. The error bars depict the standard error of the mean of three independent measurements. The peak is reached at 10 nM HER2, trastuzumab and pertuzumab respectively. The dashed line is a numeric fit with the cooperativity parameter α held constant at 1, the cyan line is a numeric fit with α freely fit.**(C)** Cryo-EM structure of pertuzumab-HER2-trastuzumab (pdb-ID 6OGE). pertuzumab (cyan) and trastuzumab (purple) bind separate epitopes on HER2 (magenta). Note that only the Fab-fragments of trastuzumab/pertuzumab were used for this structure(18).**(D)** Dissociation rates of trastuzumab (purple)/ pertuzumab (cyan) from the binary/ternary complexes. Complexes were preformed at saturating/maximal conditions (10 nM of each component). Data was fit with single-exponential dissociation models (see methods).

Titration of unlabeled HER2 to fixed concentrations of labeled trastuzumab and pertuzumab produced a characteristic hook-shaped response (Figure 3A,B). At low HER2 concentrations, increasing HER2 promoted formation of ternary complexes and increased antibody-antibody cross-correlation. At high HER2 concentrations, the cross-correlation signal decreased again, consistent with sequestration of the two antibodies by separate HER2 molecules and consequent reduction in ternary complex formation. To quantify cooperative ternary complex formation, FCCS hook titrations were analyzed using a thermodynamic equilibrium model incorporating the independently measured binary HER2 binding affinities of trastuzumab and pertuzumab (see methods). The model only accurately reproduced the observed hook-shaped dependence when allowing for a cooperativity parameter α (Figure 3B). Fitting yielded α = 3.1 ± 0.3, indicating that formation of the trastuzumab–HER2–pertuzumab ternary complex is favored approximately threefold relative to the expectation for independent binding.

We next compared dissociation kinetics of antibodies bound within binary and ternary complexes. In both cases, dissociation from ternary complexes occurred more slowly than from the corresponding binary complexes (Figure 3D). We measure 1.5-fold (pertuzumab) and 1.9-fold (trastuzumab) reduced dissociation rates for antibodies from preformed ternary complexes, suggesting that engagement of HER2 by trastuzumab and pertuzumab kinetically stabilizes the assembled complex.

## Discussion

Here we report equilibrium and kinetic characterization of trastuzumab and pertuzumab binding to HER2 in solution and directly quantify formation of the trastuzumab–HER2–pertuzumab ternary complex using fluorescence cross-correlation spectroscopy. While ternary complex formation is broadly consistent with previous structural and biophysical studies demonstrating simultaneous occupancy of distinct HER2 epitopes and little evidence for strong equilibrium cooperativity (18, 26, 29, 33), quantitative analysis reveals modest but reproducible stabilization of the assembled complex. Equilibrium measurements indicate approximately three-fold positive cooperativity in ternary complex formation, while dissociation measurements independently show a 1.5–2-fold increase in ternary complex lifetime relative to the corresponding binary interactions. Taken together, these observations provide convergent equilibrium and kinetic evidence that simultaneous engagement of HER2 by trastuzumab and pertuzumab stabilizes the assembled complex beyond that expected from strictly independent binding.

Additional insight is provided by the competitive chase experiments. When excess unlabeled antibody was added to preformed binary complexes, the reciprocal experiments revealed a clear asymmetry. Addition of pertuzumab to trastuzumab–HER2 complexes resulted in a small but reproducible reduction in the fluorescent trastuzumab signal, consistent with formation of a ternary complex accompanied by partial destabilization or redistribution of the pre-existing trastuzumab-bound population. In contrast, addition of trastuzumab to pertuzumab–HER2 complexes produced little measurable perturbation of fluorescent pertuzumab binding. Together with the slower dissociation observed for preformed ternary complexes, these findings suggest that the pathways leading to ternary complex formation are asymmetric. One possible explanation is that trastuzumab binding alters the conformational or dynamic landscape of HER2 in a manner that modestly favors subsequent pertuzumab binding, consistent with previous computational and cell-based studies proposing long-range communication between the two epitopes (32, 34). Alternatively, the observations may reflect multiple bound substates that differ slightly in stability or accessibility. Although the present data do not distinguish between these possibilities, they indicate that the ternary complex is better described as a dynamic assembly than as the simple product of two independent binding events.

These findings help clarify a long-standing discrepancy in the HER2 antibody literature. Clinical efficacy and computational studies have suggested cooperative interactions between trastuzumab and pertuzumab (12, 22, 23), whereas structural and surface-based biophysical studies established simultaneous HER2 occupancy but generally found little evidence for meaningful cooperative binding (18, 29). Our data suggests that this discrepancy may arise, at least in part, from the molecular specificity of the measurement. FCCS selectively reports on the ternary trastuzumab– HER2–pertuzumab complex through antibody–antibody cross-correlation, rather than measuring total receptor occupancy across a mixture of binary and ternary states. This true ternary readout reveals modest stabilization that may be obscured in ensemble-averaged surface measurements. At the same time, our results should not be interpreted as directly validating any single structural mechanism. Previous computational and structural studies often used Fab fragments (12, 18, 26, 33), whereas our FCCS measurements use full-length IgGs in solution. The stabilization measured here therefore reflects the net behavior of the therapeutically relevant assembly, potentially including Fab–HER2 interactions, full-IgG geometry, hinge flexibility, and local rebinding effects (28, 34).

The implications of this study extend beyond the HER2 system investigated here. Even limited evidence for cooperative interactions between trastuzumab and pertuzumab has already inspired the development of next-generation HER2-targeting therapeutics, including biparatopic antibodies such as zanidatamab that simultaneously engage multiple epitopes on the same receptor (35, 36). More broadly, many emerging therapeutic modalities—including multispecific biologics, molecular glues, targeted protein degraders, and other induced-proximity therapeutics—derive their activity from the formation and persistence of higher-order protein assemblies rather than from a single binary interaction (8–11). Rational exploitation of such mechanisms therefore requires experimental approaches capable of directly quantifying the abundance, stability, and dynamics of the molecular assemblies responsible for biological activity.

Early proponents of FCS envisioned fluctuation-based measurements as a potentially transformative approach for quantitative analysis of biomolecular interactions and pharmaceutical discovery (37, 38). While FCS and FCCS subsequently became powerful tools in mechanistic biophysics, their impact on therapeutic screening and drug-development more generally remained much more limited than initially anticipated, owing in part to instrumentation complexity and the historical emphasis on binary binding measurements and their affinities. The emergence of induced-proximity therapeutics may now alter this landscape. In such systems, the most mechanistically relevant observables are no longer affinity alone, but the abundance, cooperativity, and lifetime of higher-order molecular assemblies.

Importantly, the experiments described here were performed in a microplate-compatible format on a commercially available FCCS platform capable of automated data acquisition, demonstrating that quantitative analysis of higher-order protein assemblies is increasingly accessible beyond specialized custom-built instrumentation. This combination of robustness, automation, and direct molecular specificity may substantially broaden the practical utility of FCCS in both basic research and drug discovery. Notably, many of the opportunities envisioned in the early development of fluorescence correlation spectroscopy emerged at a time when the necessary instrumentation and therapeutic questions were not yet fully aligned. As drug discovery increasingly moves beyond binary interactions toward emergent behaviors arising from multi-component molecular assemblies, methods capable of directly interrogating the assembled species may provide insights that cannot be obtained from measurements of the individual components alone.

## Materials and Methods

### Protein preparation and fluorescent labelling

Human HER2 extracellular domain, trastuzumab, and pertuzumab (MedChemExpress, HY-P73094, HY-P9907, HY-P9912) were prepared for fluorescent labeling by exchanging the buffer into phosphate-buffered saline (PBS) at pH 7.4. Proteins were fluorescently labelled using NHS-ester derivatives of Alexa Fluor dyes (Thermo Fisher Scientific).

All proteins were incubated with a ten-fold molar excess of Alexa Fluor 546 or Alexa Fluor 647 NHS ester for 1.5 h at room temperature. Excess free dye was removed by two sequential spin desalting columns (Zeba, Thermo Scientific, 89882). Finally, the protein concentration and the degree of labelling were determined by UV–VIS spectroscopy (see supplementary Figure S2 and supplementary Table S1). Labelled proteins were stored in PBS at 4°C until use.

### Affinity titrations

All experiments were performed in glass-bottom 96-well plates (Ibidi, 89607). 12-point concentration titrations were obtained by mixing 1 nM Alexa Fluor 647 labeled trastuzumab/pertuzumab dilutions with serial dilutions of Alexa Fluor 546 labelled HER2. The final concentrations of Alexa Fluor 647 trastuzumab/pertuzumab were 0.5 nM, Alexa Fluor 546 HER2 was titrated from 40 nM to 30 pM. All experiments were performed in PBS supplemented with 2 mg/mL BSA (Sigma, A2153). A total of 80 µL of sample per well was prepared, mixed and allowed to equilibrate for 20 minutes at room temperature prior to measurement.

Ternary-complex measurements were performed equivalently to the affinity titrations. Briefly, Alexa Fluor 647 labeled trastuzumab and Alexa Fluor 546 labeled pertuzumab were mixed at fixed concentrations of 10 nM. Increasing concentrations of unlabeled HER2 (0.7 – 240 nM) were added to drive ternary-complex formation. All data points shown were prepared in triplicate.

### Dissociation kinetics of binary and ternary complexes

Dissociation kinetics of binary and ternary complexes were monitored by time-resolved FCCS. First complexes were preformed under saturating conditions. Specifically, 80 µL samples containing 10 nM Alexa Fluor 647 trastuzumab/pertuzumab were incubated with 10 nM HER2 for binary complexes, 10 nM Alexa Fluor 647 labeled trastuzumab, 10 nM Alexa Fluor 546 labeled pertuzumab and 10 nM unlabeled HER2 were incubated for ternary complexes. After 20 minutes incubation at room temperature unlabeled trastuzumab, pertuzumab or the non-specific control (NIST IgG) were added to a total concentration of 300 nM. Data was acquired periodically every 30 minutes for a total of 50 hours. Every condition was prepared in triplicate.

### FCCS data acquisition

All data was acquired using the EI-Flex Pro HT (Exciting Instruments), a well plate capable high-throughput confocal fluorescence spectrometer. All measurements lasted 60s per well using 520 nm and a 638 nm lasers at 40 µW focused in solution (20 µm above the surface of the well bottom). In addition to each described sample, confocal detection volumes were calibrated using standard fluorophores with known diffusion coefficients prior to all experiments. Namely, ATTO 655 (ATTO-TEC) and Cy3B (Lumiprobe) were used. Furthermore, spectral bleed-through was quantified using Alexa Fluor 546 HER2 samples.

### FCCS data analysis

Data analysis was performed using PhotonFit (Exciting Instruments) as well as in-house custom scripts. Autocorrelation and cross-correlation functions were calculated for all measurements and fit using a single-component diffusion model:

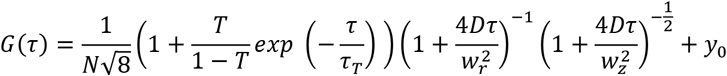

Where N is the number of molecules present in the confocal volume – the inverse of the correlation amplitude scaled by the geometric factor 1/√8. T is the triplet component amplitude, τ_T_ the triplet lifetime. D is the diffusion coefficient, and τ the lag-time. w_r_ and w_z_ are the axial and radial confocal volume waists, respectively, and y_0_ is a y-axis offset.

Replicate wells were analyzed independently and resulting fit parameters were averaged. For calibration measurements using ATTO 655 and Cy3B D was fixed to D_ATTO 655_ = 426 µm^2^s^-1^ and D_Cy3B_ = 440 µm^2^s^-1^ respectively, in accordance with previous studies (39, 40). In all other fits, w_r_ and w_z_ were fixed to the values determined by the calibration measurements. Cross-correlation curves were globally fit within titration datasets with a shared diffusion coefficient while the triplet-state contribution was fixed to zero, since triplet-state kinetics of the two dyes are independent and not expected to cross-correlate. For time course measurements the diffusion coefficients of the cross-correlation curves were globally fit for the first 10 datapoints (5 hours) and held constant for the remaining data to prevent erroneous fitting as signal decreases.

The measured cross-correlation amplitudes were corrected for the presence of uncorrelated background as well as spectral bleed-through as previously described (41, 42). The cross-correlation ratio was calculated from the ratio of N as determined from the first autocorrelation channel (520 nm excitation) and the corrected cross-correlation N.

### Affinity analysis

Antibody–HER2 binary affinities were determined by fitting the corrected relative cross-correlation data with a Hill binding model:

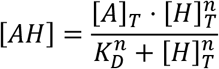

where [A]_T_ is the total labelled antibody concentration, [H]_T_ is the total HER2 concentration, K_D_ is the apparent dissociation constant (the HER2 concentration at half-maximal binding), and n is the Hill coefficient. The relative FCCS amplitude was fit as

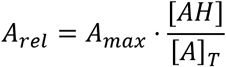

with K_D_, n and A_max_ as fitted parameters, where A_max_ is the maximal attainable relative cross-correlation amplitude. [A]_T_ was held constant at the experimental labelled antibody concentration (0.5 nM). Three replicate titrations were fit independently and the resulting parameters averaged; reported values are the mean ± standard error of the mean (n = 3 replicate titrations). See supplementary table S2 for all fitting parameters.

### Analysis of HER2 ternary complex formation

FCCS measurements of trastuzumab–HER2–pertuzumab ternary complex formation were first analyzed as for the binary reactions described above. The data were then fit to a thermodynamic equilibrium model as follows. Trastuzumab (T) and pertuzumab (P) bind to distinct epitopes on HER2 (H) with independently determined binary dissociation constants (K_T_) and (K_P_), obtained from binary FCCS affinity titrations. Binary complex formation was described by

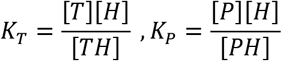

To account for potential cooperativity between the two antibodies, ternary complex formation was described using a cooperativity parameter α:

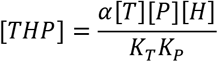

Where α values of 1 correspond to independent binding, α values greater than 1 indicate enhanced ternary complex formation, i.e. positive cooperativity, and α values less than 1 indicate reduced ternary complex formation, i.e. negative cooperativity.

Equilibrium concentrations of all species were calculated from the mass-balance constraints:

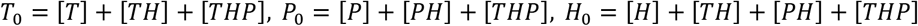

Note that the binary term [TP] denoting potential direct interaction between trastuzumab and pertuzumab was omitted from this model, since our method yields no measurable interaction (supplementary Figure S4,S5). The relative and bleed-through corrected FCCS cross-correlation signal (see above, here referred to as S) was therefore assumed to be proportional to the concentration of ternary complex and was modeled as

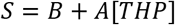

where A is a proportionality constant and B accounts for residual background signal. The cooperativity parameter and signal scaling factors were then determined by fitting the complete thermodynamic equilibrium model to the FCCS hook titration. For each iteration of the optimization, equilibrium concentrations were calculated numerically for all HER2 concentrations in the titration series, generating a predicted FCCS curve that was compared with the experimental data, see supplementary section 3 for details on the implementation and supplementary table 3 for all fit parameters.

## Acknowledgements

The authors thank Joseph Allcock and Fraser Davies for software development and support and Elliot Steele, Jonathan Walsh and Damith Vithanage for instrument development and support.

## Author contributions

S.H.M., M.A.S.A. and T.D.C conceptualized research; S.H.M. designed the study; S.H.M. and G.M. performed research; S.H.M. and M.A.S.A. analyzed data; S.H.M., M.A.S.A., C.P.T. and T.D.C. wrote the manuscript.

