## Supplementory Information for "Fluorescence cross-correlation spectroscopy quantifies affinity, cooperativity, and kinetic stability in ternary protein complexes"

### Supplementary Methods

#### 1. Protein labelling and dye characterization

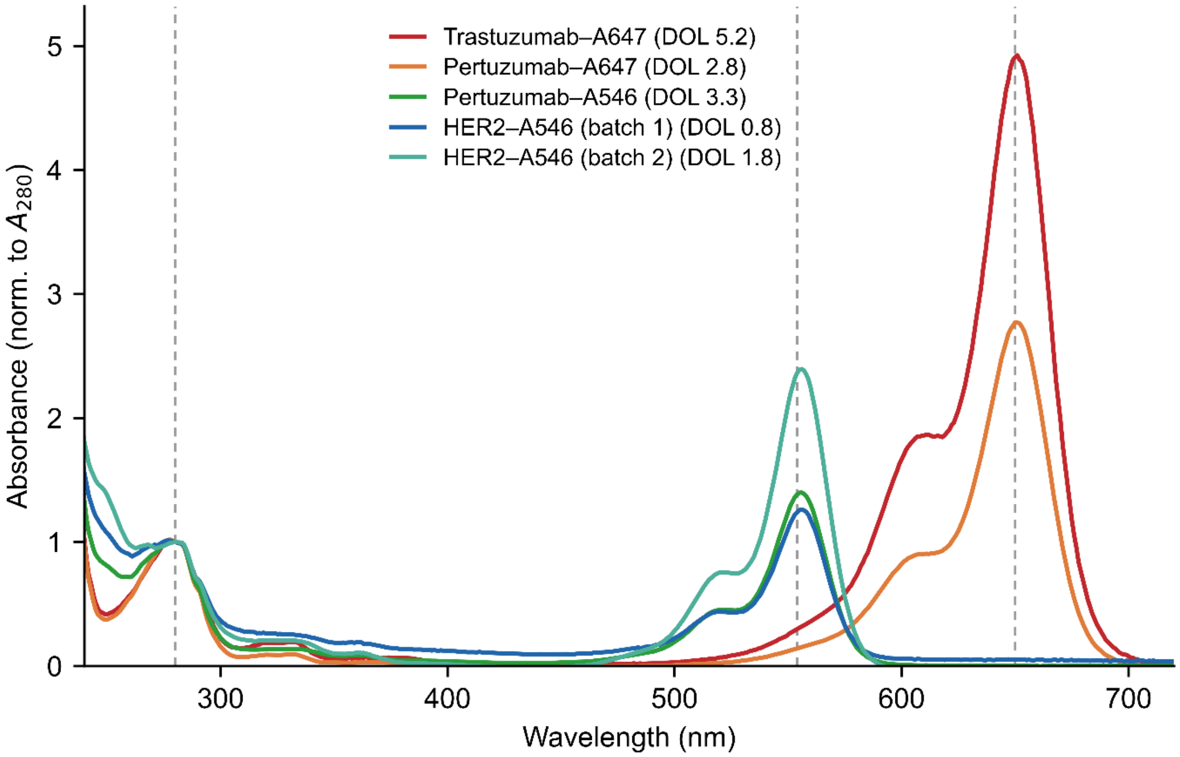


***Figure S1.*** *UV–Vis absorbance spectra of Alexa Fluor–labelled antibody and HER2 conjugates, normalized to the protein absorbance at 280 nm. Dashed vertical lines mark 280 nm and the dye absorbance maxima at 554 nm (AF546) and 650 nm (AF647). The degree of labelling (DOL) of each conjugate is given in the legend.*

| Sample | Dye | $\varepsilon_{280}$ ($M^{-1}cm^{-1}$) | $A_{280}$ | A_dye | [Protein] (μM) | [Dye] (μM) | DOL |
|---|---|---|---|---|---|---|---|
| Trastuzumab–A647 | AF647 | 215,000 | 0.151 | 0.739 | 5.97 | 30.92 | 5.18 |
| Pertuzumab–A647 | AF647 | 222,000 | 0.245 | 0.678 | 10.11 | 28.37 | 2.81 |
| Pertuzumab–A546 | AF546 | 222,000 | 0.246 | 0.341 | 9.25 | 30.43 | 3.29 |
| HER2–A546 (batch 1) | AF546 | 62,340 | 0.088 | 0.109 | 1.20 | 0.98 | 0.81 |
| HER2–A546 (batch 2) | AF546 | 62,340 | 0.178 | 0.422 | 2.05 | 3.77 | 1.84 |

***Table S1.*** *Degree of labelling (DOL) of antibody and HER2 conjugates determined by UV–Vis absorbance. Protein concentrations are stock values (dilution-corrected); absorbances are buffer-subtracted and measured at 1 cm pathlength. AF546 and AF647 denote Alexa Fluor 546 and 647;* $\varepsilon_{280}$ *is the protein molar extinction coefficient at 280 nm.*

#### 2. Binary affinity titrations

| Antibody | Replicate | KD (nM) | Amax | $R^2$ | RMSE | n |
|---|---|---|---|---|---|---|
| **Trastuzumab** | 1 | 1.44 ± 0.33 | 0.199 ± 0.010 | 0.963 | 0.014 | 12 |
| | 2 | 1.51 ± 0.22 | 0.205 ± 0.007 | 0.985 | 0.009 | 12 |
| | 3 | 1.07 ± 0.34 | 0.198 ± 0.013 | 0.931 | 0.019 | 12 |
| | mean ± s.e.m. | 1.34 ± 0.14 | 0.201 ± 0.002 | 0.959 | — | 3 |
| **Pertuzumab** | 1 | 1.25 ± 0.36 | 0.376 ± 0.023 | 0.939 | 0.033 | 12 |
| | 2 | 1.32 ± 0.31 | 0.378 ± 0.019 | 0.961 | 0.027 | 12 |
| | 3 | 0.99 ± 0.26 | 0.340 ± 0.018 | 0.950 | 0.027 | 12 |
| | mean ± s.e.m. | 1.18 ± 0.10 | 0.365 ± 0.012 | 0.950 | — | 3 |

**Table S2. Binary-affinity FCCS fits.** *Dissociation constants (KD) and saturation amplitudes (Amax) from the 1:1 depletion-aware model (Eq. S1–S2), fit unweighted to each replicate titration (12 HER2 concentrations, 0–40 nM; [A]T = 0.5 nM fixed). Parameter uncertainties for individual replicates are the fit standard errors; the mean* *row is mean ± s.e.m. over the three replicates. $R^2$ and RMSE are on the corrected relative cross-correlation amplitude.*

#### 3. Thermodynamic model of ternary complex formation

Formation of trastuzumab–HER2–pertuzumab ternary complexes was analyzed using a thermodynamic equilibrium model based on the law of mass action. The model considers six molecular species: free trastuzumab (T), free pertuzumab (P), free HER2 (H), the binary complexes trastuzumab–HER2 (TH) and pertuzumab–HER2 (PH), and the ternary complex trastuzumab–HER2–pertuzumab (THP).

Binary binding is described by the independently measured dissociation constants

$$K_T = \frac{[T][H]}{[\mathrm{TH}]}$$

and

$$K_P = \frac{[P][H]}{[\mathrm{PH}]}.$$

To account for cooperative ternary complex formation, the equilibrium concentration of the ternary complex was expressed as

$$[\mathrm{THP}] = \frac{\alpha[T][P][H]}{K_T K_P},$$

where $\alpha$ is the cooperativity factor. In this formulation,

- $\alpha = 1$ corresponds to independent binding,
- $\alpha > 1$ indicates positive cooperativity,
- $\alpha < 1$ indicates negative cooperativity.

Accordingly, the effective dissociation constants for the second binding event become

$$K_{P|TH} = \frac{K_P}{\alpha}$$

and

$$K_{T|PH} = \frac{K_T}{\alpha}.$$

Thus, positive cooperativity corresponds to an increased affinity of the second antibody once the first antibody is already bound.

**Mass-balance equations**

The total concentrations of trastuzumab, pertuzumab and HER2 satisfy

$$T_0 = [T] + [\mathrm{TH}] + [\mathrm{THP}],$$

$$P_0 = [P] + [\mathrm{PH}] + [\mathrm{THP}],$$

$$H_0 = [H] + [\mathrm{TH}] + [\mathrm{PH}] + [\mathrm{THP}].$$

Substituting the mass-action expressions gives

$$T_0 = [T] + \frac{[T][H]}{K_T} + \frac{\alpha[T][P][H]}{K_T K_P},$$

$$P_0 = [P] + \frac{[P][H]}{K_P} + \frac{\alpha[T][P][H]}{K_T K_P},$$

$$H_0 = [H] + \frac{[T][H]}{K_T} + \frac{[P][H]}{K_P} + \frac{\alpha[T][P][H]}{K_T K_P}.$$

These equations constitute three coupled nonlinear equations in the three unknown equilibrium concentrations $[T]$, $[P]$, $[H]$. Once the free concentrations have been determined, the concentrations of all binary and ternary complexes follow directly from the equilibrium relationships above.

**Numerical solution**

No analytical solution of the coupled equilibrium equations was used. Instead, equilibrium concentrations were obtained numerically for every experimental HER2 concentration. For each HER2 concentration, the residual vector

$$r = \begin{pmatrix} T_0 - [T] - [\mathrm{TH}] - [\mathrm{THP}] \\ P_0 - [P] - [\mathrm{PH}] - [\mathrm{THP}] \\ H_0 - [H] - [\mathrm{TH}] - [\mathrm{PH}] - [\mathrm{THP}] \end{pmatrix}$$

was minimized subject to the physical constraints

$$[T] \geq 0,\ \ [P] \geq 0,\ \ [H] \geq 0.$$

Optimization was performed using a bounded nonlinear least-squares algorithm until the mass-balance equations were simultaneously satisfied. The resulting free concentrations were subsequently used to calculate the concentrations of all binary and ternary species.

This numerical procedure yields the exact thermodynamic equilibrium solution without requiring an explicit analytical expression for ternary complex formation and readily accommodates cooperative interactions.

**Prediction of FCCS measurements**

The experimentally measured FCCS cross-correlation signal was assumed to be proportional to the equilibrium concentration of ternary complex,

$$S = B + A[\mathrm{THP}],$$

where $A$ is an arbitrary scaling constant relating ternary complex concentration to the measured FCCS signal, and $B$ accounts for residual background cross-correlation and systematic offsets.

This observation model allows the measured FCCS amplitudes to be compared directly with the thermodynamic equilibrium predictions without requiring absolute calibration of the signal.

**Parameter estimation**

Binary dissociation constants were fixed to the values independently determined from binary FCCS affinity titrations,

$$K_T = 1.34 \text{ nM},$$

$$K_P = 1.18 \text{ nM}.$$

The fitted parameters therefore comprised $A,\ B,\ \alpha.$

To ensure positivity and improve numerical stability, cooperativity was optimized in logarithmic space,

$$x = \log_{10}(\alpha),$$

with

$$= 10^x,$$

where $x$ is the fitted optimization parameter.

For a given trial parameter set, the complete equilibrium model was solved numerically at every HER2 concentration in the titration series, generating a predicted FCCS hook curve. Parameter estimation was performed by nonlinear weighted least-squares minimization of the residuals between the predicted and experimentally measured FCCS signals over the complete titration series.

**Statistical analysis**

Goodness of fit was assessed using the residual sum of squares (RSS), the coefficient of determination ($R^2$), and the Akaike Information Criterion (AIC). Parameter uncertainties were estimated from the covariance matrix returned by the nonlinear least-squares optimization. Standard errors were obtained from the square roots of the diagonal elements of the covariance matrix. Ninety-five percent confidence intervals for the cooperativity factor were calculated by propagation of the uncertainty in the fitted logarithmic parameter,

$$\alpha = 10^x,$$

which naturally yields asymmetric confidence intervals in linear space. The fitted cooperativity factor was reported together with its corresponding confidence interval as the principal quantitative measure of cooperative ternary complex formation.

Fits used total antibody concentrations $T_0 = P_0 = 10$ nM and were weighted by the per-point standard error of the mean (n = 12 HER2 concentrations, three replicates each). The binary dissociation constants $K_T$ and $K_P$ were held fixed at the binary-titration values; models in which $K_T$ and $K_P$ were fitted freely were degenerate ($K_P$ collapsed to its lower bound), which is why the dissociation constants were fixed. Uncertainties are ±1 SE; the 95% confidence interval for α was obtained by covariance propagation and confirmed by profile likelihood. The cooperative model is strongly preferred (ΔAIC = −32.4).

| **Parameter** | **Independent model (α = 1)** | **Cooperative model** |
|---|---|---|
| Fixed dissociation constants | $K_T = 1.34$ nM, $K_P = 1.18$ nM | $K_T = 1.34$ nM, $K_P = 1.18$ nM |
| No. of free parameters, k | 2 | 3 |
| Amplitude, A | 0.0386 ± 0.0006 | 0.0277 ± 0.0008 |
| Background, B | 0.0022 ± 0.0009 | 0.0040 ± 0.0009 |
| Cooperativity, α | 1 (fixed) | 3.17 ± 0.34 |
| α, 95% CI | — | 2.57 – 3.91 |
| $R^2$ | 0.865 | 0.979 |
| Residual sum of squares (RSS) | $4.30 \times 10^{-3}$ | $6.70 \times 10^{-4}$ |
| Reduced $\chi^2$ | 11.8 | 0.75 |
| AIC | 31.4 | −1.0 |
| ΔAIC vs independent | 0 (reference) | −32.4 |

***Table S3.*** *Fitted parameters, goodness-of-fit statistics and model comparison for the numerical thermodynamic fits of the HER2 ternary hook titration.*

#### 4. Supplementary Controls

**Confocal-volume calibration**

The detection volume of each channel was calibrated from free-dye reference solutions of known diffusion coefficient: Cy3B (green channel, D = 440 $\mu m^2/s$) and Atto655 (red channel, D = 426 $\mu m^2/s$)(1, 2). Autocorrelations were fitted with D held fixed, leaving the lateral and axial radii $w_r$ and $w_z$ free. Typical confocal volumes were $V_{eff}$ = 0.53 ± 0.06 fL (green) and 0.56 ± 0.06 fL (red). These values were stable throughout the run ($w_r$ CV ≤ 1.7 %, $V_{eff}$ CV ≈ 11 %, n = 100), and the per-molecule brightness of both dyes was constant to within ~9 % CV. This calibration was carried out daily before any other measurements were performed.

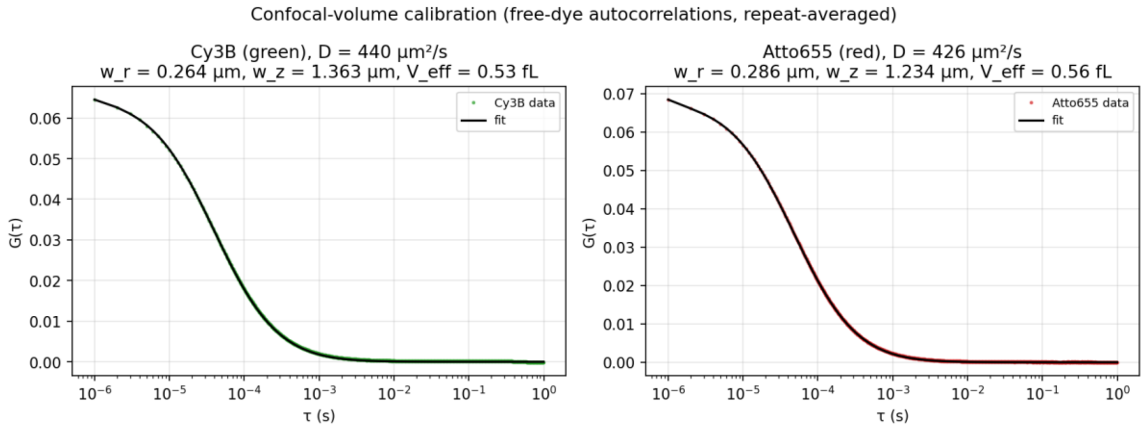


**Figure S2**. Confocal-volume calibration. Repeat-averaged autocorrelations of the free dyes Cy3B (green channel) and Atto655 (red channel), fitted with the diffusion coefficient fixed to the literature value to extract the detection-volume dimensions (black lines, fits).

***Spectral bleed-through***

Bleed-through was quantified from a singly-labelled (Alexa Fluor 542) HER2 control well as the ratio of background-corrected red to green count rates, $k_{gr} = (F_{red} - BG_{red})/(F_{green} - BG_{green})$, where green refers to the signal emitted by Alexa Fluor 542 labeled species and red to the signal emitted by Alexa Fluor 647 labeled species. The green-labelled control produced 524 ± 20 kHz in the green channel and 25.9 ± 1.0 kHz in the red channel, giving a bleed-through fraction $k_{gr}$ = 0.048 ± 0.0003 (4.8 %), stable to within 0.5 % CV over 50 h. Red-to-green bleed-through, assessed from the red-only (Alexa Fluor 647-labelled trastuzumab) control wells, was < 1 % of the red signal (green count rate 4–5 kHz at red count rates of 470–760 kHz). Cross-correlation amplitudes were corrected for green-to-red bleed-through using the measured $k_{gr}$.

***Background counts.***

Constant background offsets of $BG_{green}$ = 3.9 kHz and $BG_{red}$ = 1.0 kHz were applied in the bleed-through and cross-correlation corrections. The green background was taken from the residual green-channel signal of the red-only control wells; the red background was set to a nominal 1.0 kHz, as no dedicated buffer-only blank was included in this plate. Both backgrounds were small relative to the specific sample count rates (typically < 1 %) and were stable over the run.

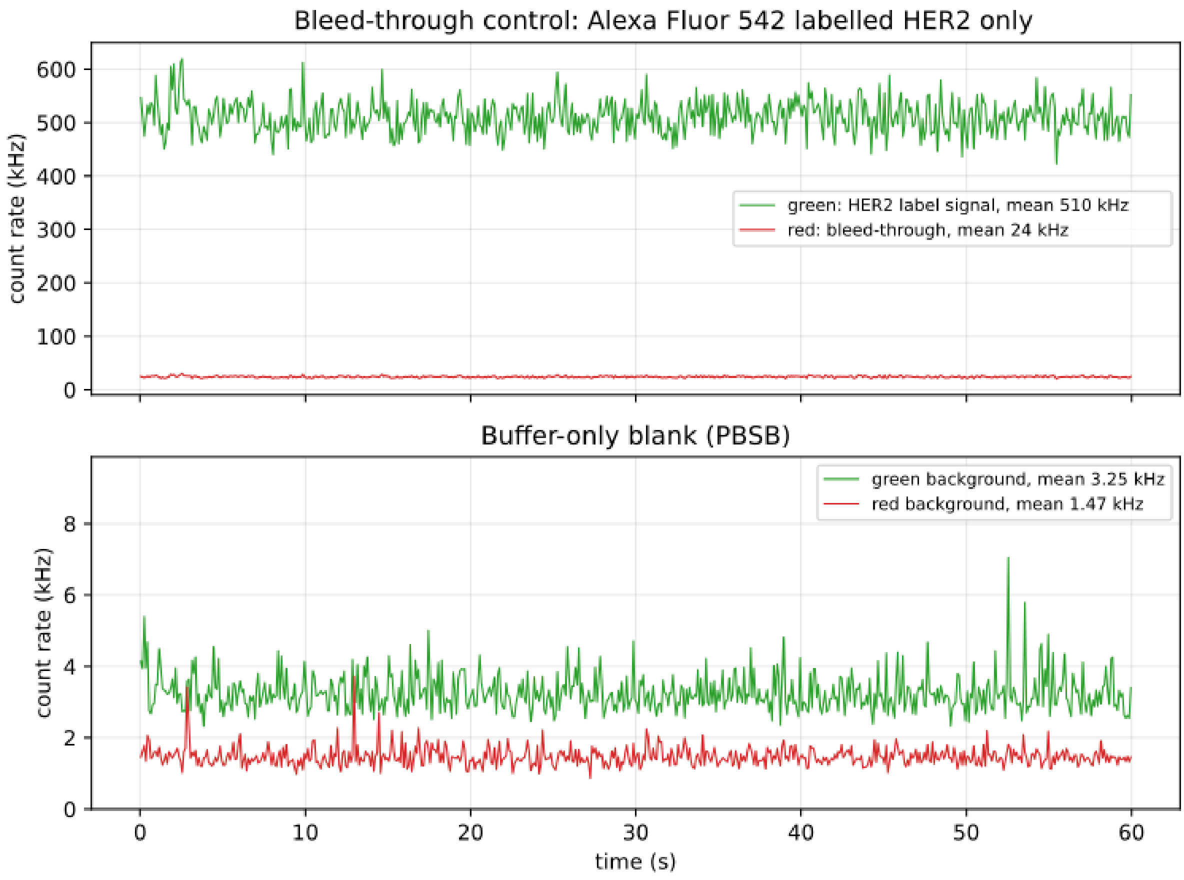


***Figure S3***. *Bleed-through and background. Top: photon count-rate trace of a Alexa Fluor 542 labeled HER2 control; the red-channel signal (24 kHz) relative to the green signal (510 kHz) gives the green→red bleed-through (k_gr ≈ 4.7 %). Bottom: buffer-only (PBSB) blank, giving the green (3.3 kHz) and red (1.5 kHz) background count rates.*

##### 3.2 No-cross-correlation controls

To confirm that the FCCS cross-correlation reports specifically on HER2-bridged co-diffusion of the two antibodies — rather than on spectral bleed-through, antibody–antibody association, or optical artefacts — a mixture of labelled trastuzumab and labelled pertuzumab (one green-, one red-labelled) was measured in the absence and presence of HER2 under identical conditions. For each sample the green and red autocorrelations and the green–red cross-correlation were fitted (Figures S2, S3), and the cross-correlation amplitude was converted to a bleed-through- and background-corrected cross-correlation ratio (Figure S4).

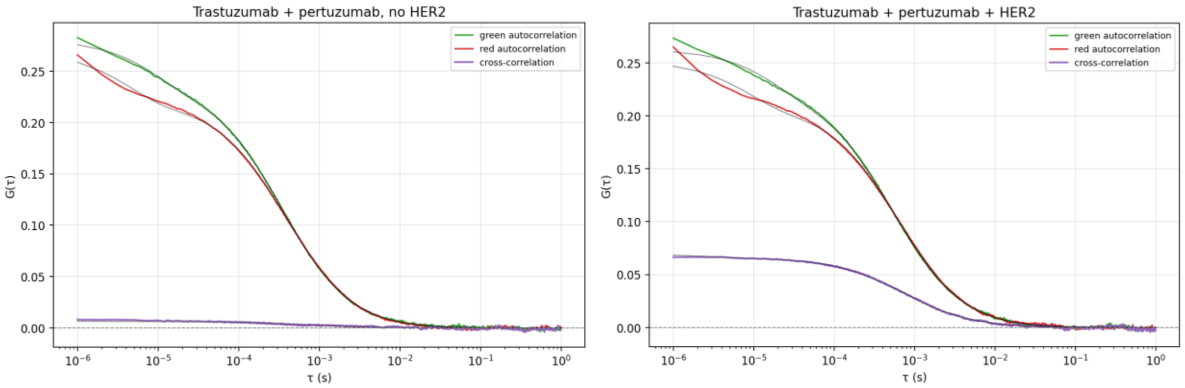


***Figure S4***. *Left: Trastuzumab + pertuzumab without HER2. Green and red autocorrelations are large while the green–red cross-correlation (purple) is mostly flat. Note, the small amplitude can be attributed to spectral bleed-through and is later corrected for. Right: Trastuzumab + pertuzumab + HER2. Clear increase in cross-correlation (purple) relative to the two auto-correlations in green and red.*

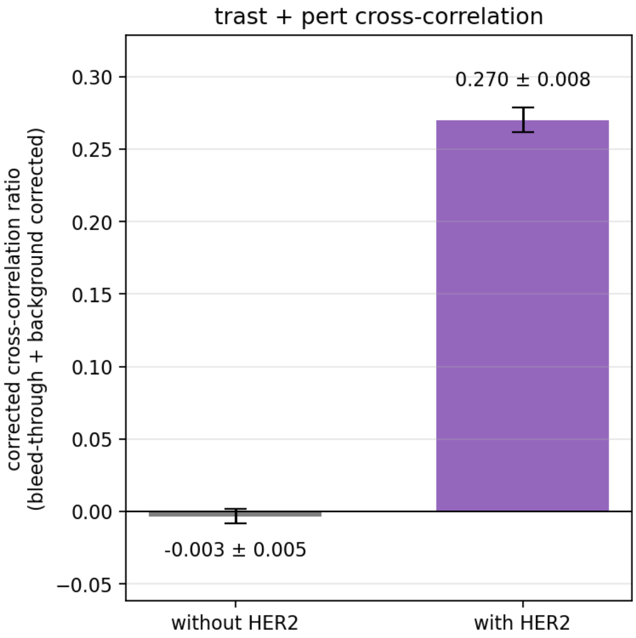


***Figure S5***. *Bleed-through- and background-corrected cross-correlation ratio of trastuzumab+ pertuzumab without (grey) and with HER2 (purple, mean ± SEM over n = 10 time-segmented sub-acquisitions, with the uncertainty in the bleed-through fraction propagated into the error bars). The ratio is zero within error without HER2 (−0.003 ± 0.005) and rises to 0.27 ± 0.01 with HER2, confirming HER2-specific ternary complex formation.*

#### Dissociation kinetics

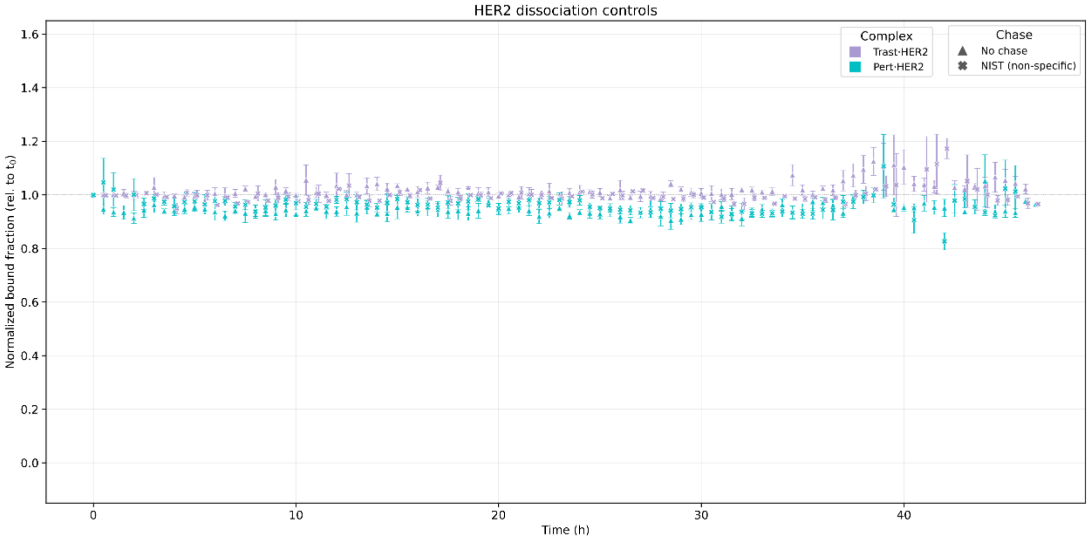


***Figure S6.*** *Stability of trastuzumab-HER2 (purple) and pertuzumab-HER2(cyan) complexes when chased with non-specific NIST-IgG (X) or not chased at all (triangles). The cross-correlation ratio stays approximately stable over 48h.*
